# Effects of the COVID-19 pandemic on publication landscape in chimeric antigen receptor-modified immune cell research

**DOI:** 10.1101/2021.06.01.446639

**Authors:** Ahmet Yilmaz, Jianhua Yu

**Affiliations:** The Ohio State University, Comprehensive Cancer Center, Columbus, OH 43210, USA; Department of Hematology and Hematopoietic Cell Transplantation, City of Hope National Medical Center, 1500 E. Duarte Road, KCRB, Bldg. 158, Room 3017, Los Angeles, CA 91010, USA; Hematologic Malignancies Research Institute, City of Hope National Medical Center, Los Angeles, CA 91010, USA; Department of Immuno-Oncology, City of Hope Beckman Research Institute, Los Angeles, CA 91010, USA; City of Hope Comprehensive Cancer Center and Beckman Research Institute, Los Angeles, CA 91010, USA

**Keywords:** Chimeric antigen receptor, T cells, NK cells, Cancer immunotherapy, Coronavirus

## Abstract

Chimeric antigen receptors (CARs) are artificial receptors introduced mainly into T cells. CAR-induced immune cell (CARi) products have achieved impressive success rates in treating some difficult-to-treat hematological malignancies. Here, we describe effects of the global COVID-19 pandemic on CARi publication landscape. Due to the pandemic, the total number of publications decreased in 2020 compared to 2019 in all fields of cancer immunotherapy except CARi. Nearly exponential increases in the number of CARi publications slowed-down in 2020 for the first time in the past 11 years. There were more CARi than coronavirus publications until 2020 when coronavirus publications increased over 5,000% compared to 2019 (575 publications in 2019 vs. 30,390 in 2020). Unlike cancer immunotherapy where the majority of the publications consist of conference abstracts and review articles, majority of the coronavirus publications are original research articles. There are more coronavirus publications in Pubmed than Embase. The opposite is true for CARi publications. Our analysis of the data from the FDA Adverse Event Reporting System (FAERS) show significantly higher death rate in patients treated with Kymirah than Yescarta (28.14% vs. 16.02%). Kymirah and Yescarta are the two main CAR T cell products for treatment of DLBCL and/or B-ALL. However, despite being highly significant, this result is not easily interpretable due to multiple confounding variables in the FAERS data. Our analysis additionally suggest that the significant effects of co-stimulatory domains (4-1BB vs. CD28) consistently reported in preclinical studies do not translate into clinical results. Our manual curation of the CARi publications in PubMed shows that only 5.2% of the publications report results from CARi clinical trials, although we found 663 clinical trials listed on ClinicalTrials.gov database. In conclusion, publication landscape in CARi as well as other fields of cancer immunotherapy has changed due to the global COVID-19 pandemic. This trend will likely continue in the near future. CARi research is now in need of increased measures by publishers to reduce repetitive and/or duplicate publications and more stringent criteria for data entry into public databases including PubMed, Embase, ClinicalTrials.gov, and FAERS to advance this important field of medical research.

## The COVID-19 pandemic has re-shaped the publication landscape in chimeric antigen receptor-modified immune cell research

The global COVID-19 pandemic is probably one of the most significant natural disasters in modern times with major biological, economical, and social impacts on the society at large. Effects of the pandemic on publication landscape in cancer immunotherapy research have not been thoroughly investigated in the literature. Here, we provide a comprehensive overview of the changes in the field due to the pandemic with a focus on chimeric antigen receptor (CAR)-modified immune cell (CARi) research.

CARs are artificial receptors introduced mainly into T cells^1^. We and others have shown that CAR-induced natural killer (CAR NK) cells are likely as effective and offer additional advantages compared to the CAR-induced T (CAR T) cells^2–6^. CARs are composed of an extracellular singlechain variable fragment (scFv) domain derived from the heavy and light chains of monoclonal antibodies raised against tumor-associated antigens (TAA) attached to intracellular activating domain(s) such as CD3ζ and/or co-stimulatory domain(s) such as CD28 or 4-1BB^6^. The binding of the extracellular domain to the TAA, which may be as specific as in binding by monoclonal antibodies, activates the intracellular domain(s), which in turn activates the immune cell, resulting in the killing of tumor cells.

CARi research has been one of the most exciting fields in immunotherapy. Anti-CD19 CAR T cells have achieved unprecedented success rates as high as 90% in treating relapsed or refractory acute lymphoblastic leukemia (ALL) and other B cell leukemia and lymphomas, bringing fresh hope to cancer patients and prompting the FDA to issue breakthrough designations^7^. CARi therapy is one of the costliest types of immunotherapy, but it is covered by The Centers for Medicare and Medicaid Services in the U.S. The CARi technology is projected to create a market in the range of billions of U.S. dollars in the near future^8^. Along with this huge clinical success has come a boom in the number of publications.

Publications containing the terms “chimeric antigen receptor” in the title or abstract have ballooned from a single publication in 1989 to 1,391 in 2020 in PubMed (Medline) and 1,629 in Embase (Excerpta Medica dataBASE) (Fig. 1A). As of January 2021, there are 4,192 and 7,140 CARi publications in PubMed and Embase, respectively. Moreover, 2,725 publications are shared between the two databases whereas 348 and 4,035 publications are unique to the Medline and Embase, respectively (Fig. 1B). Each year since 2010 has seen a nearly exponential increase in the number of CARi publications except that in 2020, a year heavily affected by COVID-19, where the total number of publications in Embase seemed to reach a plateau (Fig. 1A); the publication growth rate was only 4.78% compared to 2019 (1,555 vs. 1,632), the lowest in the past 11 years. This slow-down in CARi publications was not seen in PubMed likely because PubMed includes no conference abstracts and more review manuscripts than Embase.

**Fig. 1.**
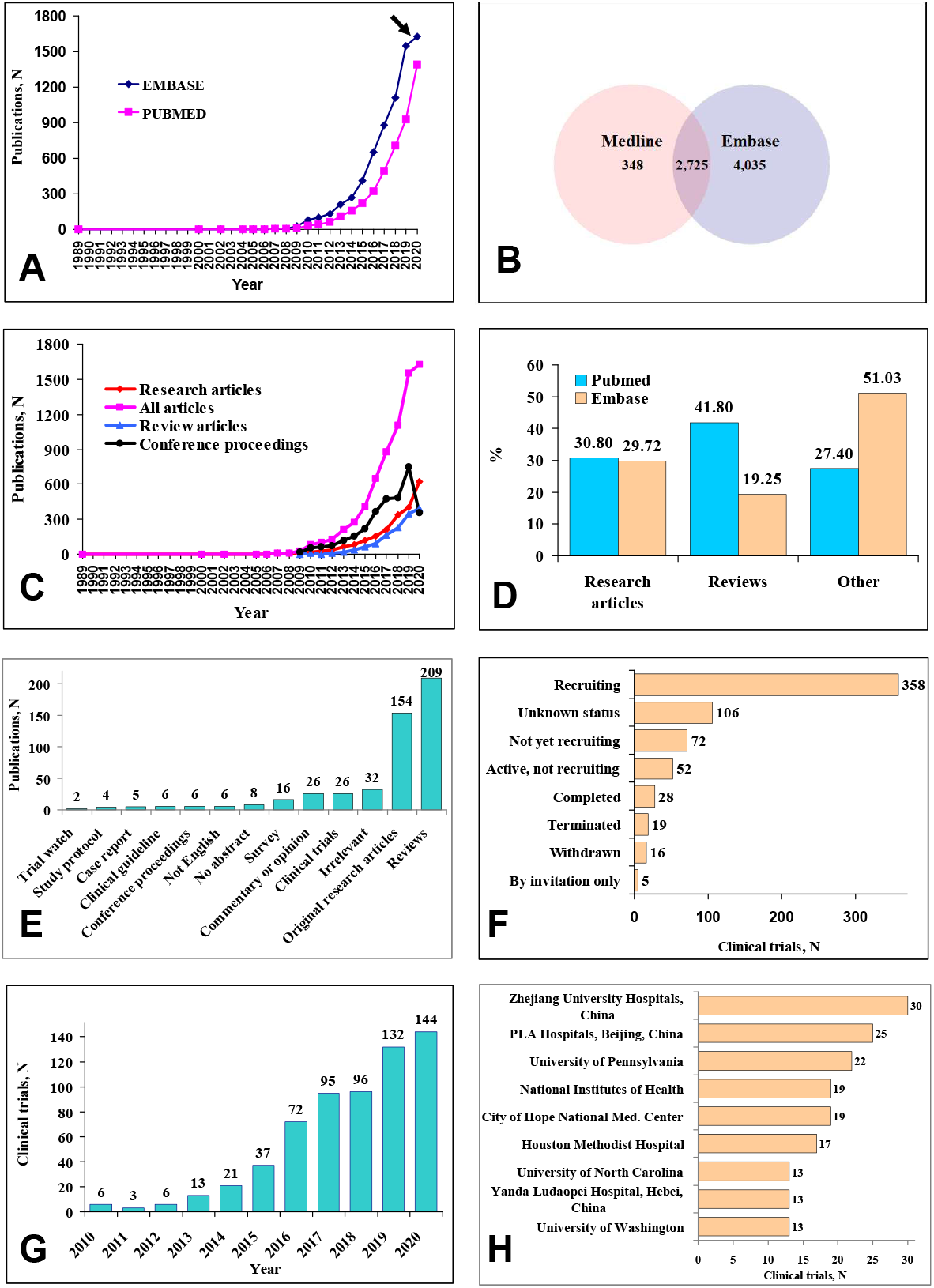
*A* CAR-modified immune cell (CARi) publications in Embase and PubMed by year of publication. *B* Total CARi publications from 1989 to 2021 shared between PubMed and Embase. *C* CARi publication types in Embase. *D* Percent publication types in Embase vs. PubMed. *E* Types of CARi publications in PubMed. *F* Status of CARi clinical trials as of 2021. *G* CARi clinical trials conducted per year. Trials without a clear start date or those that are projected to start in the future are not included, *H* Notable institutions conducting CARi clinical trials.

Plotting CARi publication types in Embase by year revealed a sharp decrease in the number of conference abstracts (746 in 2019 vs. 374 in 2020) and a clear slow-down in the number of review articles (351 in 2019 vs. 404 in 2020), although original research articles grew at a rate similar to the previous years (411 in 2019 vs. 771 in 2020, Fig. 1C). We manually curated 500 CARi publications in PubMed and compared the types of publications in PubMed and Embase (Figs. 1D and 1E). Despite the similar percent research articles (30.8% vs. 29.72%), there is nearly twice in number more review articles (41.8% vs. 19.25%) and significantly less “other” (i.e., publications that are neither original research articles nor reviews) type of manuscripts in PubMed than Embase (27.4% vs. 51.03%, Fig. 1D).

Interestingly, only 5.2% of the publications (26 of the 500 representative publications corresponding to 205 of the 3,937 publications in total) in PubMed report results from CARi clinical trials, although we found 663 clinical trials listed on *ClinicalTrials.gov* when we searched the terms “chimeric antigen receptor, chimeric antigen receptor T, chimeric antigen receptor NK, chimeric antigen receptor natural killer, CAR, CAR T, CAR NK, and CAR natural killer” and removed duplicates and trials missing critical information such as targets or disease types (Figs. 1F and 1G, Supp. Figs. 1–2, Supp. Table 1). Unlike preclinical studies that focused on B cell markers as well as solid tumors, the clinical trials mostly targeted B cell markers alone or in combination with antibodies or modified or multiple CARs (Supp. Tables 2–5).

The largest number of clinical trials has been registered by (with number of trials conducted in parentheses) Zhejiang University (30) and People’s Liberation Army General Hospitals (25) in China and University of Pennsylvania (22), City of Hope National Medical Center, and National Institutes of Health (19 each) in the U.S. (Fig. 1H, Supp. Table 6). These five institutions conduct 17.4% of all clinical trials worldwide. In total, 56.2% and 36.8% of all clinical trials are being conducted in China and U.S., respectively; only 7% of the clinical trials are being conducted in the rest of the world.

Although the top three Chinese institutions conduct more trials than the top three U.S. institutions (68 vs. 60, Fig. 1H), the latter institutions published 2.2-fold (772 vs. 348) more total manuscripts including 2.4-fold (220 vs. 92) more review articles than their Chinese counterparts as revealed by searching the terms “Zhejiang”, “Beijing”, “Hebei”, “Pennsylvania”, “City of Hope”, or “National Institutes of Health” in PubMed along with the terms “chimeric antigen receptor”. Interestingly, Zhejiang University Hospitals, a Chinese institution conducting the largest number of clinical trials, conducted more trials (30 vs. 22) but published 4.1-fold less (599 vs. 140) manuscripts than its American counterpart, University of Pennsylvania. Although limited by possible language barriers, these results suggest that Chinese institutions conduct more clinical trials but publish fewer manuscripts than U.S. institutions.

In order to investigate if the slow-down in CARi publications in Embase was associated with the global COVID-19 pandemic or changes in other types of cancer immunotherapy, we compared the number of publications in CARi, coronavirus, antibody-drug conjugates (ADC), anti-PD-1 (pembrolizumab), and anti-CTLA4 (ipilimumab) monoclonal antibody research (Figs. 2–5). Inhibitors of PD-1 and CTLA-4, the two checkpoint blockers, along with ADC represent the two other major fields of research in cancer immunotherapy in addition to CARi. There was a higher percent of review articles in CARi than in the other cancer immunotherapy fields. The total number of publications decreased in 2020, compared to 2019, in all fields of immunotherapy except CARi. In contrast, coronavirus publications spiked at unprecedented proportions (5,185%, 575 in 2019 vs. 30,390 in 2020).

**Fig. 2.**
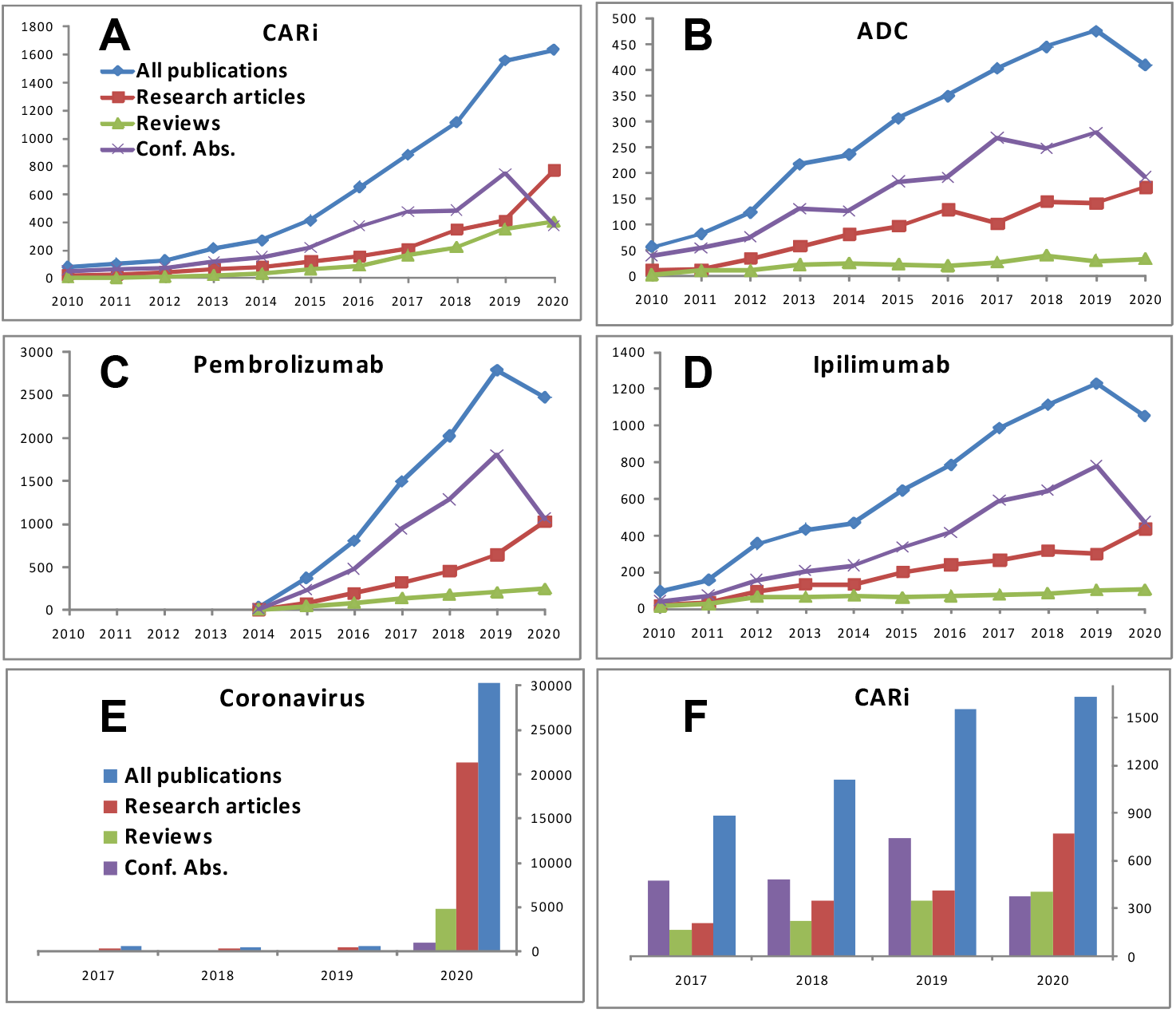
Publications in coronavirus and cancer immunotherapy research plotted by year. Total number of publications decreased in all cancer immunotherapy fields examined except CARi (A - D). When all publications are considered, there are nearly twice more research articles and less review articles and conference abstracts in coronavirus than CARi research (E - F). X axis = year of publication, Y axis = number of publications. The legend in A applies to A – D, and the legend in E applies to E and F. Data are current as of February, 2021. Conf. Abs. = conference abstracts, All publications = all publication types combined.

**Figure 3.**
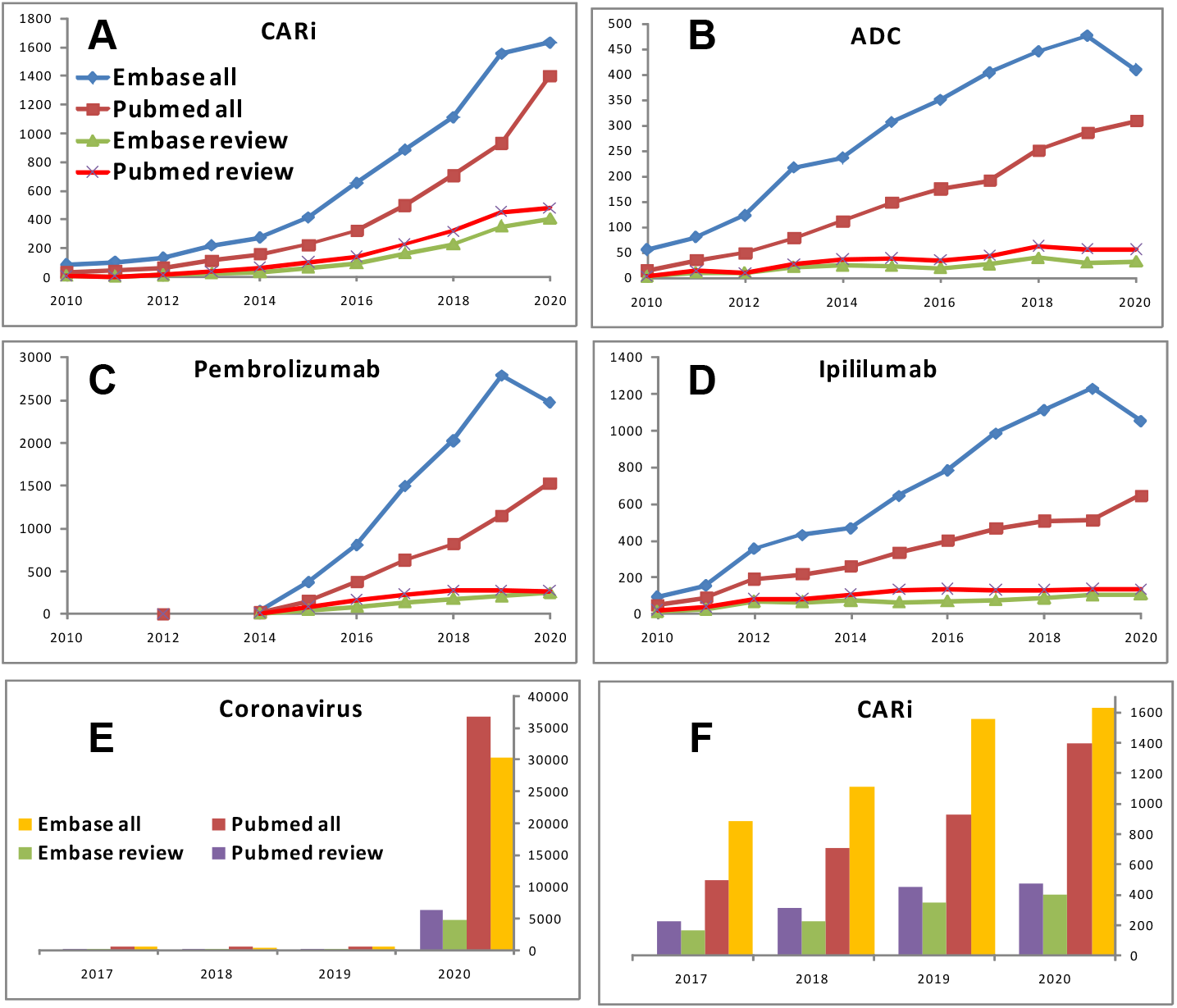
A comparison of cancer immunotherapy and coronavirus publications in Embase vs. PubMed from 2010 to 2020. When all publication types are considered, there are more cancer immunotherapy publications in Embase than PubMed (A - D). There is an opposite trend in coronavirus research where there are more publications in PubMed than Embase (E - F). Reviews as a percentage of all publications is lower in coronavirus than CARi research in both PubMed and Embase. X axis = year of publication, Y axis = number of publications. The legend in A applies to the Figures A - D, and the legend in E applies to Figures E and F. All = all publication types, including reviews, original research articles, conference abstracts, etc.

**Figure 4.**
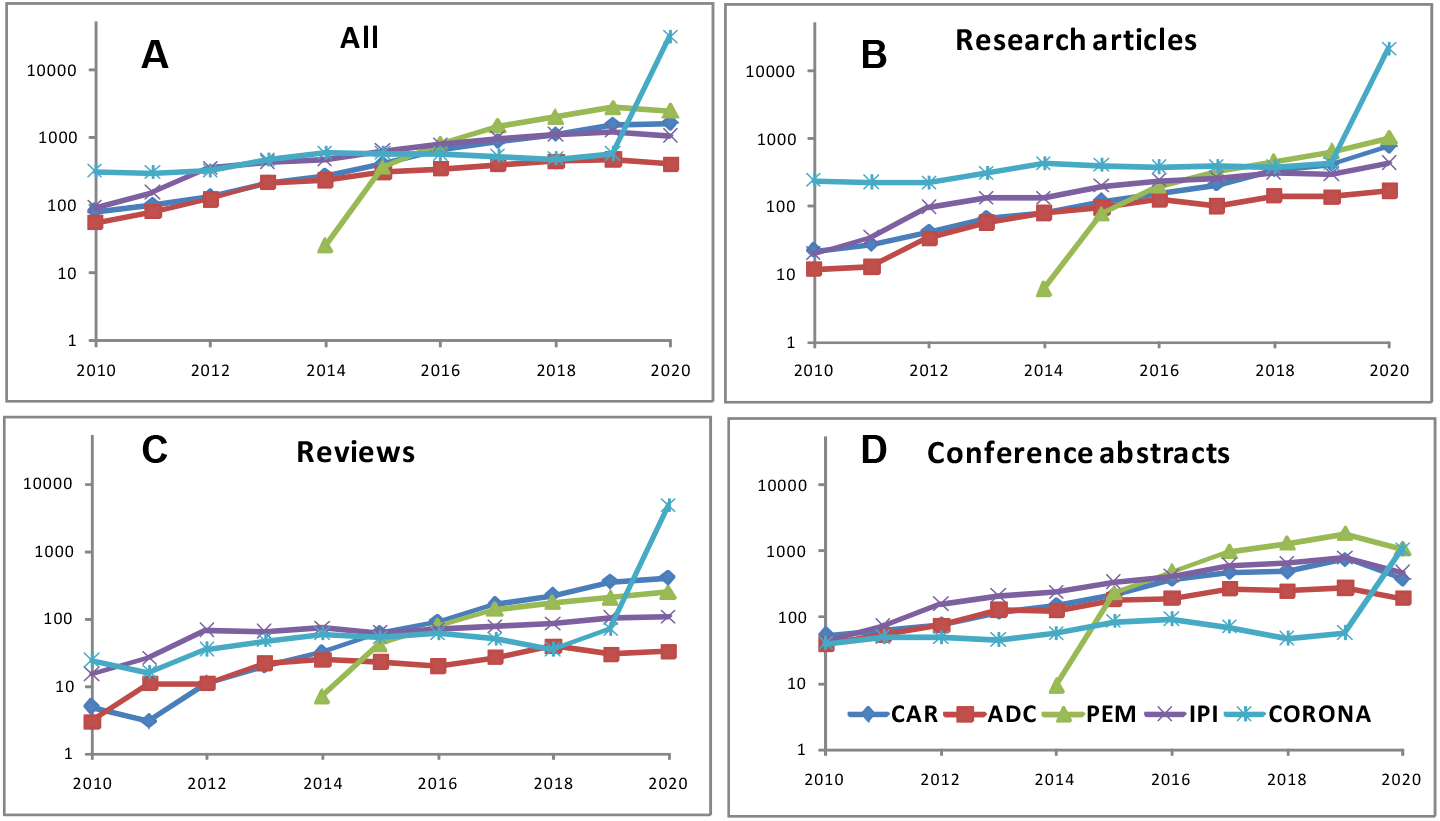
A comparison of publication types in cancer immunotherapy and coronavirus research from 2010 to 2020. Historically, there have been significantly more research articles published in coronavirus than cancer immunotherapy research. Unlike coronavirus publications with a steady growth rate over the years, cancer immunotherapy publications quickly grew and surpassed the coronavirus publications beginning in 2018. However, in the year 2020 when the pandemic took over, there were nearly 10 times more coronavirus publications than all other cancer immunotherapy fields combined. Growth in the number of conference abstracts is especially striking in pembrolizumab (PEM) research. The number of PEM conference abstracts was the largest among all fields only two years after its first publication in 2014 and have remained the largest ever since. The number of conference abstracts was lowest in coronavirus than in any other field but, in 2020, they dramatically increased and reached a level slightly less than PEM (1,034 vs. 1,066). The Y axis is drawn to the log10 scale. All = all publication types including original research articles, reviews, conference abstracts, etc; X axis = year of publication; Y axis = number of publications. The legend in Figure D applies to all graphs. CAR = chimeric antigen receptor modified immune cells, PEM = pembrolizumab, CORONA = coronavirus, IPI = ipilimumab, ADC = antibody drug conjugate.

**Figure 5.**
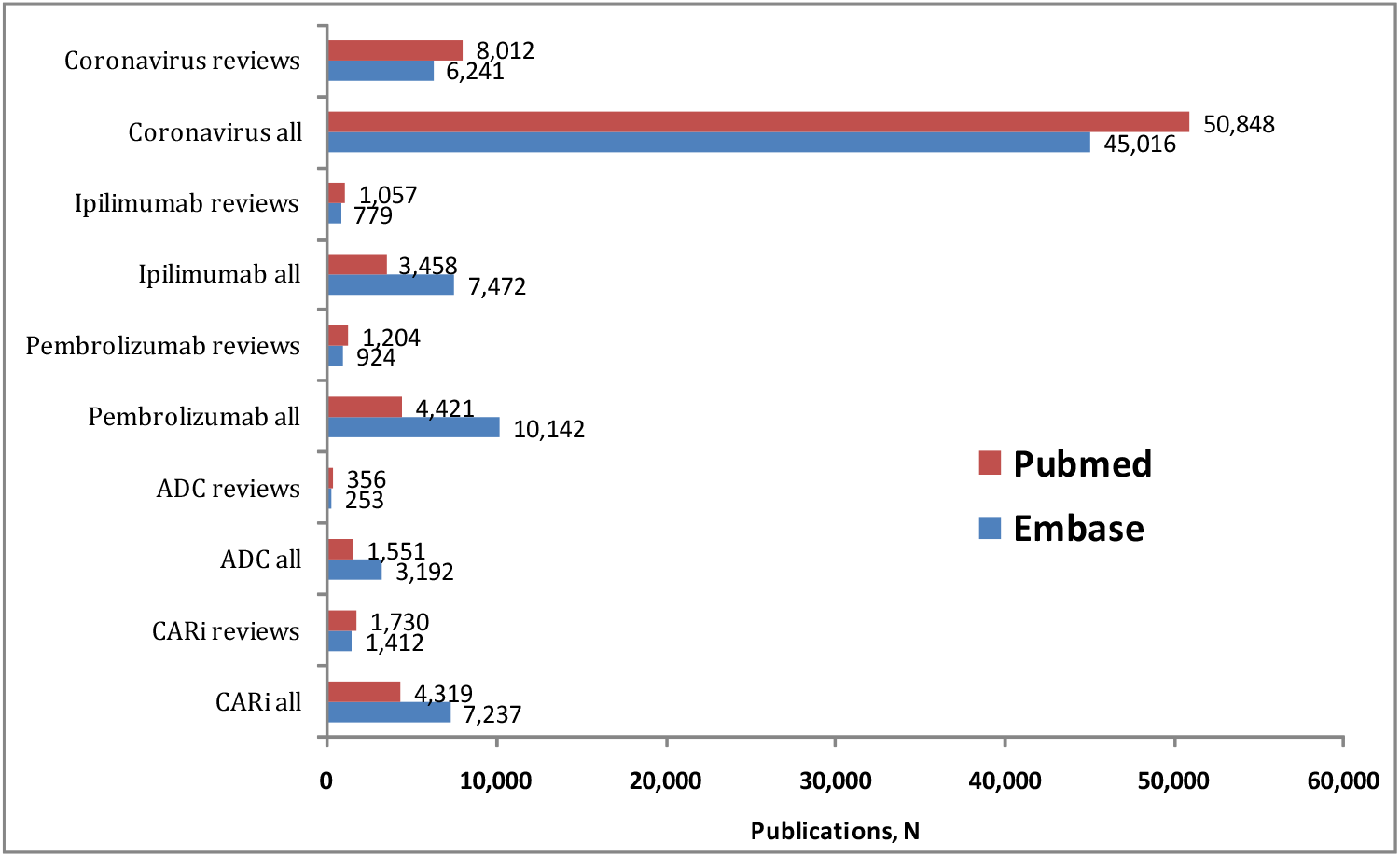
Total cancer immunotherapy and coronavirus publications from 1989 to 2020 in PubMed vs. Embase. PubMed contains fewer total but more review articles than Embase in all cancer immunotherapy fields. In coronavirus research, however, there are more both overall and review manuscripts in PubMed than Embase. All = all publication types including original research articles, reviews, conference abstracts, etc; reviews = review manuscripts; CARi = chimeric antigen receptor-modified immune cells; ADC = antibody drug conjugate.

Unlike cancer immunotherapy where majority of the publications consist of conference abstracts and review articles, most of the coronavirus publications were original research articles. In 2020, there was a higher percent of review articles (24.75% vs 15.71%) and conference abstracts (22.92% vs. 3.4%) but less original research articles (47.24% vs. 70.3%) published in CARi than coronavirus research. Conference abstracts decreased in all immunotherapy fields in 2020 but spiked in coronavirus research. These results suggest an overall slow-down or decrease in the number of publications in cancer immunotherapy research publications coinciding with the global COVID-19 pandemic.

Our manual curation of the CARi publications in PubMed revealed that some potentially high-impact CARi research questions have yet to be satisfactorily answered in the literature. For example, studies investigating CARi target selection, the first critical step in creating novel CARi cell products, through search of gene expression databases such as Gene Expression Omnibus by The National Center for Biotechnology Information (NCBI), The Cancer Genome Atlas (TCGA), or the Expression Atlas by European Molecular Biology Laboratory (EMBL) to discover highly and uniformly expressed genes in cancer subtypes or research focusing on CARi signaling, have rarely been explored. On the contrary, redundant, duplicate, or minimally relevant publications have increased at an alarming rate. The most frequent objective in original research articles involves trying to optimize CAR T cell design.

Safety of CAR T cell products is investigated in 12.5% of the sampled publications in PubMed. The four CAR T cell products already approved by the U.S. FDA include Kymirah (tisagenlecleucel) by Novartis, Breyanzi (lisocabtagene maraleucel) by Juno Therapeutics (now a part of Celgene), and Yescarta (axicabtagene ciloleucel) and Tecartus (brexucabtagene autoleucel) by Kite Pharma (now a part of Gilead Sciences) (Supp. Table 7). Kymirah and Yescarta were approved for B-ALL and diffuse large B-cell lymphoma (DLBCL), respectively, in 2017. Kymirah and Brenyazi were approved for DLBCL in 2018 and 2021, respectively. Tecartus was approved for mantle cell lymphoma in 2020.

FDA has created a public database, FDA Adverse Event Reporting System (FAERS) where adverse events (AEs) can be entered by consumers, manufacturers, or health care professionals mainly for post-marketing surveillance. As of January 2021, only three AEs including one death in patients treated with Tecartus have been entered into the database (Supp. Table 7), likely due to its recent approval. Further, 2,072 AEs in patients treated with Yescarta have been reported; 1,972 of these events were serious, and 332 (16.02%) resulted in death. As for Kymirah, 1,546 AEs have been reported, where 1,461 of which were serious and 435 (28.14%) resulted in death. Although 50 AEs, all of which were serious, including 9 deaths (18%), have been reported in patients treated with Breyanzi, all 50 events occurred before the approval of Breyanzi by the FDA in February 2021. Therefore, they may not be considered as “post-marketing” events.

Although limited by multiple confounding variables, the higher death rate reported in patients with AEs treated with Kymirah (28%) than Yescarta (16.02%) and Breyanzi (18%) is still intriguing. Disease type could be important in these differences because Yescarta and Breyanzi are only indicated for DLBCL, whereas Kymirah is indicated for both DLBCL and B-ALL. An interesting observation can be made regarding the role of co-stimulatory domains in these death rates. Preclinical studies have consistently shown that 4-1BB is safer than CD28^9^. However, the death rate is higher in Kymirah, which contains 4-1BB, than Yescarta, which contains CD28. In addition, death rates in Breyanzi and Yescarta, which have different co-stimulatory domains but indicated for the same disease, are similar. These observations suggest that the significant effects of costimulatory domains reported in preclinical studies do not necessarily translate into real-world scenarios.

These data from FAERS, an official database, are difficult to interpret due to many confounding variables. Reporting AEs to FAERS is currently voluntary. Therefore, some severe AEs may not have been reported. It is unknown how many patients were treated successfully and, thus, were not reported. Additionally, the number of AEs in patients treated with CARi products in FAERS has an unusually wide range (1 to 35), suggesting that some institutions may have entered only the partial AE profile of patients. Several variables such as event date, age, or weight have been entered for some patients but not for others. It is not clear whether FDA has any oversight at all on data entry. In addition, these drugs are administered to different patient populations, likely following different protocols. Considering that the patients are heavily pre-treated before starting the CAR T therapy, most patients likely suffer from multiple conditions. It is not clear if or how many of these AEs are due to secondary conditions, or other drugs used to treat them, rather than the CAR T therapy itself.

All of the anti-CD19 CAR T products including Kymirah, Yescarta, Tecartus, and Breyanzi were granted “fast-track” approval by the FDA based on the JULIET (Phase II), ZUMA-1 (Phase I/II), ZUMA-2 (Phase II), and TRANSCEND-NHL-001 (Phase I) clinical trials, respectively. We are not aware of any Phase III clinical trials reported to evaluate these products. Well-controlled, randomized clinical trials are needed to assess the efficacy and safety profiles of these CAR-modified T cell products. Until then, the pivotal trials that led to the FDA approval, and were recently reviewed by us^10^ and others, are perhaps the best sources of information to evaluate the efficacy and safety of these CAR T products.

The number of CARi and coronavirus publications obtained through the “results by year” function of the new version of PubMed released in November 2019 differs from those obtained through the classical “search” method. To create a representative database allowing a detailed analysis, we entered the terms “dabrafenib lung” in PubMed in February, 2021. The search returned 130 publications. However, there were 151 publications in the timeline obtained through the “results by year” function. In order to investigate the reason for this discrepancy, we downloaded the raw Medline files directly from the NCBI databases through Entrez using scripts executed in PERL, an open source programming language, in a Unix shell using the same search terms. This strategy returned 130 publications. A detailed analysis of the 151 publications retrieved through the “results by year” approach revealed that 26 publications had different online and actual publication dates, suggesting that these 26 publications were likely entered twice in databases used to obtain the “results by year” timeline although it is unknown why the search produced 151 rather than 156 publications. We included only data obtained through the “search” function of the PubMed in this manuscript as the “results by year” timeline does not seem reliable.

In conclusion, publication landscape in CARi as well as other fields of cancer immunotherapy has changed due to the global COVID-19 pandemic. This trend will likely continue in the near future as scientists try to contain the pandemic. CARi research is now in need of increased measures by publishers to reduce non-peer-reviewed, repetitive and/or duplicate publications, at least in high-impact journals, and more stringent criteria for data entry into public databases including *PubMed, Embase, ClinicalTrials.gov*, and FAERS. These measures have the potential to significantly ease the burden on scientists trying to obtain reliable data that they urgently need to advance this important field of medical research.

## Acknowledgements

The authors regret that it was not possible to include many interesting studies in the field due to limited space.

## Authors’ contributions

All authors wrote the initial draft, revised the review, finalized the last version of the article, and approved the final version.

## Funding

This work was supported by grants from the NIH (AI129582, CA247550, NS106170, CA223400), the Leukemia & Lymphoma Society (1364-19), and The California Institute for Regenerative Medicine (DISC2COVID19-11947).

## Competing interests

Dr. Jianhua Yu is the co-founder of the Cytoimmune, Inc.

---

**Supplementary Figure 1.**
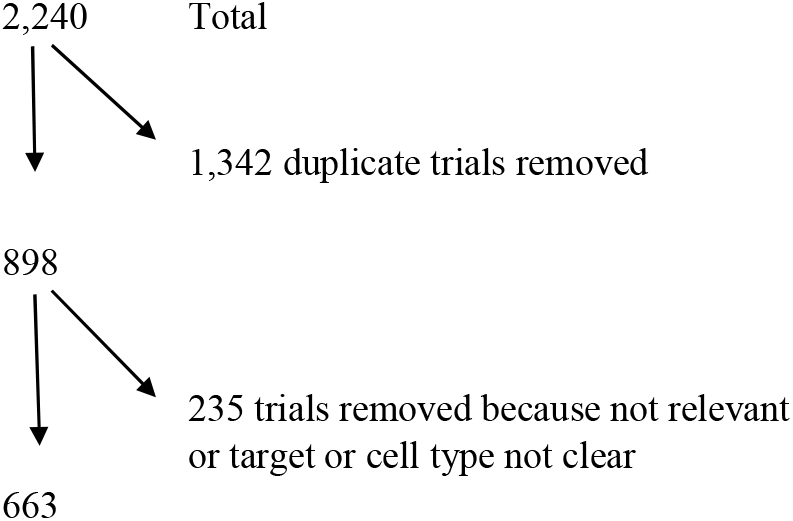
Search of the terms “chimeric antigen receptor, chimeric antigen receptor T, chimeric antigen receptor NK, chimeric antigen receptor natural killer, CAR, CAR T, CAR NK, and CAR natural killer” on *ClinicalTrials.gov* returned 2,240 clinical trials. Only 663 clinical trials remained after removal of duplicates or trials lacking critical information such as target or cell type.

**Supplementary Figure 2.**
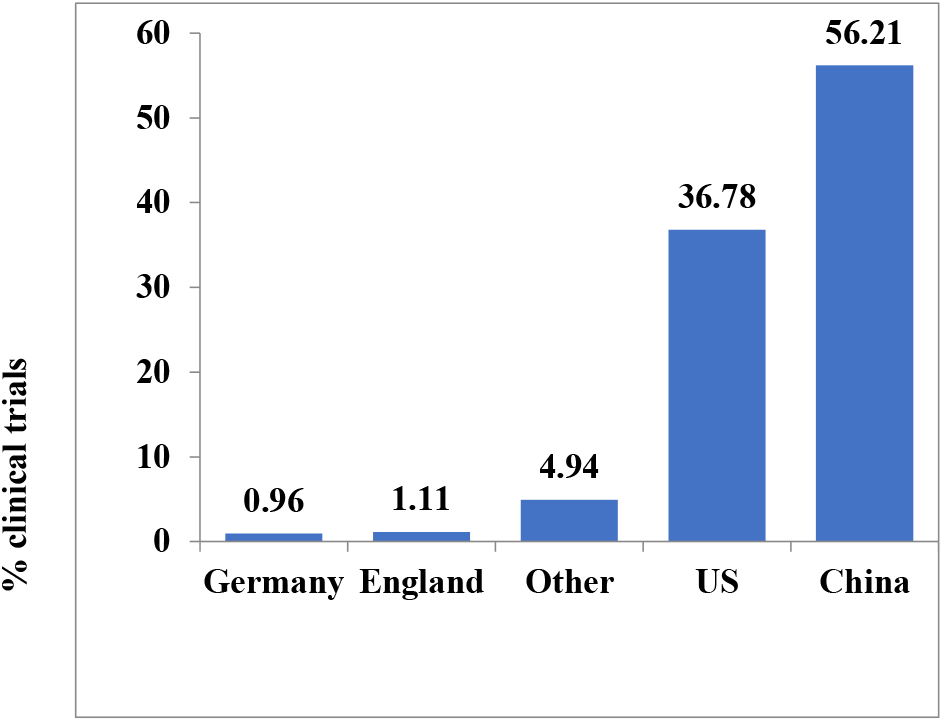
Percent clinical trials conducted by country from 2003 to 2020^a^ ^a^6, 7, 231 and 353 clinical trials are registered in Germany, England, US, and China, respectively. Twelve clinical trials are registered in other countries.

**Supplementary Table 1.** Clinical trials retrieved when the terms below were searched on *ClinicalTrials.gov* in January, 2021.

| **Search term entered** | **Clinical trials found, *N*** |
| --- | --- |
| Chimeric antigen receptor | 358 |
| Chimeric antigen receptor T | 313 |
| Chimeric antigen receptor NK | 6 |
| Chimeric antigen receptor natural killer | 6 |
| CAR | 810 |
| CAR T | 723 |
| CAR NK | 16 |
| CAR natural killer | 8 |
| **TOTAL** | **2,240** |

**Supplementary Table 2.** Tumor-associated antigens targeted in chimeric antigen receptor-modified immune cell (CARi) clinical trials. % = percent of all clinical trials; *N* = number of clinical trials.

| **Clinical trials** |  | **Target** | **Alias** | **Function of the target** |
| --- | --- | --- | --- | --- |
| *N* | % |  |  |  |
| 208 | 31.42 | CD19<sup>a</sup> | B-lymphocyte antigen | B cell development and function<sup>1</sup> |
| 69 | 10.42 | BCMA<sup>b</sup> | B-cell maturation antigen | Implicated in development of multiple myeloma<sup>2</sup>, important in survival and maturation of B cells<sup>2,3</sup>, important in autoimmunity<sup>4</sup> |
| 26 | 3.93 | Mesothelin<sup>c</sup> | Mesothelin | A tumor differentiation antigen present on the mesothelial cells<sup>5</sup> |
| 25 | 3.78 | CD19/CD22 | CD22 (SIGLEC2) | CD22 is an inhibitory co-receptor in signaling through B cell receptor<sup>6</sup>, it is highly expressed in ALL<sup>7</sup> and inhibits autoimmune disease, along with FcγIIb, by silencing B cells<sup>8</sup>. Anti-CD22 CAR T cells induce remission in B-ALL resistant to CD19 CAR T therapy<sup>9</sup> |
| 23 | 3.47 | CD19/CD20 | CD20 (MS4A1) | A pan B cell marker along with CD19, CD20 is important in B cell signaling and development<sup>10</sup>; its expression is lost upon differentiation into plasma cells, suggesting that it may be important in differentiation of B-cells into plasma cells<sup>11</sup>. In addition, it mediates T-cell independent antibody responses<sup>12</sup>. Anti-CD20 antibodies are used for treating B cell malignancies |
| 20 | 3.02 | GD2<sup>d</sup> | Disialoganglioside-2 | Highly and specifically expressed in neuronal tumors<sup>13</sup> |
| 18 | 2.72 | CD22 | SIGLEC-2 | See above |
| 17 | 2.57 | CD30 | TNFRSF8 | A Type 2 T cell marker<sup>14</sup> |
| 16 | 2.42 | CD123 | IL-3R | Highly expressed in AML subtypes; it is a marker of leukemic stem cells<sup>15</sup> |
| 16 | 2.42 | GP3 | Glypican 3 | Implicated in bone and skeletal muscle cell proliferation and differentiation<sup>16</sup> |
| 13 | 1.96 | HER2 | Receptor tyrosine-protein kinase ErbB-2 | An oncogene in the EGFR family important especially in breast cancer<sup>17</sup> |
| 10 | 1.51 | EGFR<sup>e</sup> | ErbB-1 | Along with HER2, Her 3, and Her 4, EGFR is a member of receptor tyrosine kinases important in oncogenesis and signal transduction<sup>18</sup> |
| 9 | 1.36 | BCMA | B-cell maturation antigen | See above |
| 9 | 1.36 | MUC1<sup>f</sup> | Mucin 1, epithelial membrane antigen | MUC1, overexpressed in cancerous epithelial cells lining epithelial surfaces in the lungs, intestine, etc. is a marker for solid tumors<sup>19</sup> |
| 9 | 1.36 | PSCA/PMSA | PSCA, prostate stem cell antigen / PSMA, prostate-specific membrane antigen | Increased PSCA expression is a marker for bladder, pancreatic, and prostate cancers<sup>20</sup>; PSMA, a glutamate carboxypeptidase that cleaves N-acetylaspartylglutamate, is important in diagnosis of prostate cancer<sup>21</sup> |
| 8 | 1.21 | CD7 | GP40 | Important in lymphocyte development and T and B-cell trafficking. It is targeted in T cell-acute lymphoblastic leukemia<sup>22</sup> due to its role in apoptosis in activated T lymphocytes<sup>23</sup> |
| 7 | 1.06 | CEA | Carcinoembryonic antigen | Important in signaling, cell adhesion, and metastasis<sup>24</sup> |
| 6 | 0.91 | CD33 | SIGLEC3 | Anti-CD33 monoclonal antibody conjugated with calicheamicin (gemtuzumab ozogamicin) has been approved by FDA for newly diagnosed and relapsed or refractory acute myeloid leukemia<sup>25</sup> |
| 6 | 0.91 | EFGRvIII<sup>g</sup> | ErbB-1 | A deletion mutant version of EGFR frequently found in brain tumors<sup>26</sup> |
| 6 | 0.91 | EpCAM | Epithelial cell adhesion molecule | Important in epithelial cell to cell adhesion, proliferation, and differentiation<sup>27</sup> |
| 6 | 0.91 | NKG2D | NKG2D | Recognizes stress-induced surface ligands<sup>28</sup> |
| 5 | 0.76 | B7-H3 | CD276, B7 Homolog 3 | B7-H3 is a pro-tumorigenic immune checkpoint protein and a strong inhibitor of T cell proliferation and activation<sup>29,30</sup> |
| 5 | 0.76 | CD20 | MS4A1 | See above |
| 4 | 0.6 | Claudin18.2 | Claudin18.2 | Claudin 18.2 (CLDN18.2) is an important component of the tight junctions in epithelial cells; it is highly expressed in gastric tumors<sup>31,32</sup> |
| 4 | 0.6 | ROR1/ROR2 | NTRKR1 / NTRKR2 | ROR1, important in cell migration, is a prognostic factor for triple negative breast cancer<sup>33</sup>; ROR2, important in cartilage and growth plate development<sup>34</sup>, has prognostic value in colorectal cancers<sup>35</sup> |
| 3 | 0.45 | CD38 | cyclic ADP ribose hydrolase | Anti-CD38 antibodies daratumumab and isatuximab are established treatment options for multiple myeloma<sup>22</sup> |
| 3 | 0.45 | CD70 | TNFSF7 | CD70 is a marker for T and B cell activation (i.e., absent in resting T cells but induced upon activation), and mediates suppressor functions of TGF-beta especially on effector memory T cells<sup>36</sup> |
| 3 | 0.45 | CS1 | SLAMF7 | Marker for malignant plasma cells, highly expressed in multiple myeloma<sup>37,38</sup> |
| 3 | 0.45 | IL13Rα2 | CD213A2 | Plays a role in binding and internalization of IL13, overexpressed in brain tumors<sup>39</sup> |
| 3 | 0.45 | Lewis Y | Lewis Y | Lewis Y is a carbohydrate antigen highly expressed in a range of solid tumors<sup>40</sup> |
| 3 | 0.45 | NKG2DL | NKG2DL | Ligand for the activating immunoreceptor NKG2D, important in activation of cytotoxic lymphocytes through recognition of MHC class I-related glycoproteins on virally infected or malignant cells<sup>41</sup> |
| 2 | 0.3 | CD138 | Syndecan 1 | Syndecan 1 is a transmembrane protein important in tumor cell proliferation, angiogenesis, matrix remodeling and migration<sup>42</sup> |
| 2 | 0.3 | CD147 | Basigin | Basigin is a transmembrane glycoprotein receptor important in tumor metastasis and virus entry into cells especially during SARS-CoV-2 infection<sup>43</sup> |
| 1 | 0.15 | CD171 | L1CAM | Important in cell adhesion, development of the nervous system, and prognosis in several cancers<sup>44</sup> |
| 85 | 12.84 | Others | Not applicable | Not applicable |

<sup>a</sup>CD19 alone or in combination with IL-7, CCL19, IL-15, IL-13R $\alpha$ 2, iCasp9, or acalabrutinib
<sup>b</sup>BCMA alone or in combination with CD19, BiRD, lenalidomide, CD29, CD38, CS1, or PD-1
<sup>c</sup>Mesothelin alone or in combination with PD-1 or CTLA-4
<sup>d</sup>GD2 alone or in combination with IL-7R, IL-15, iCasp9, CD276, or PSMA
<sup>e</sup>EGFR alone or in combination with PD-1, CXCR5, or IL-12
<sup>f</sup>MUC1 alone or in combination with PD-1 or CTLA-4
<sup>g</sup>EFGRvIII alone or in combination with pembrolizumab

**Supplementary Table 3.**
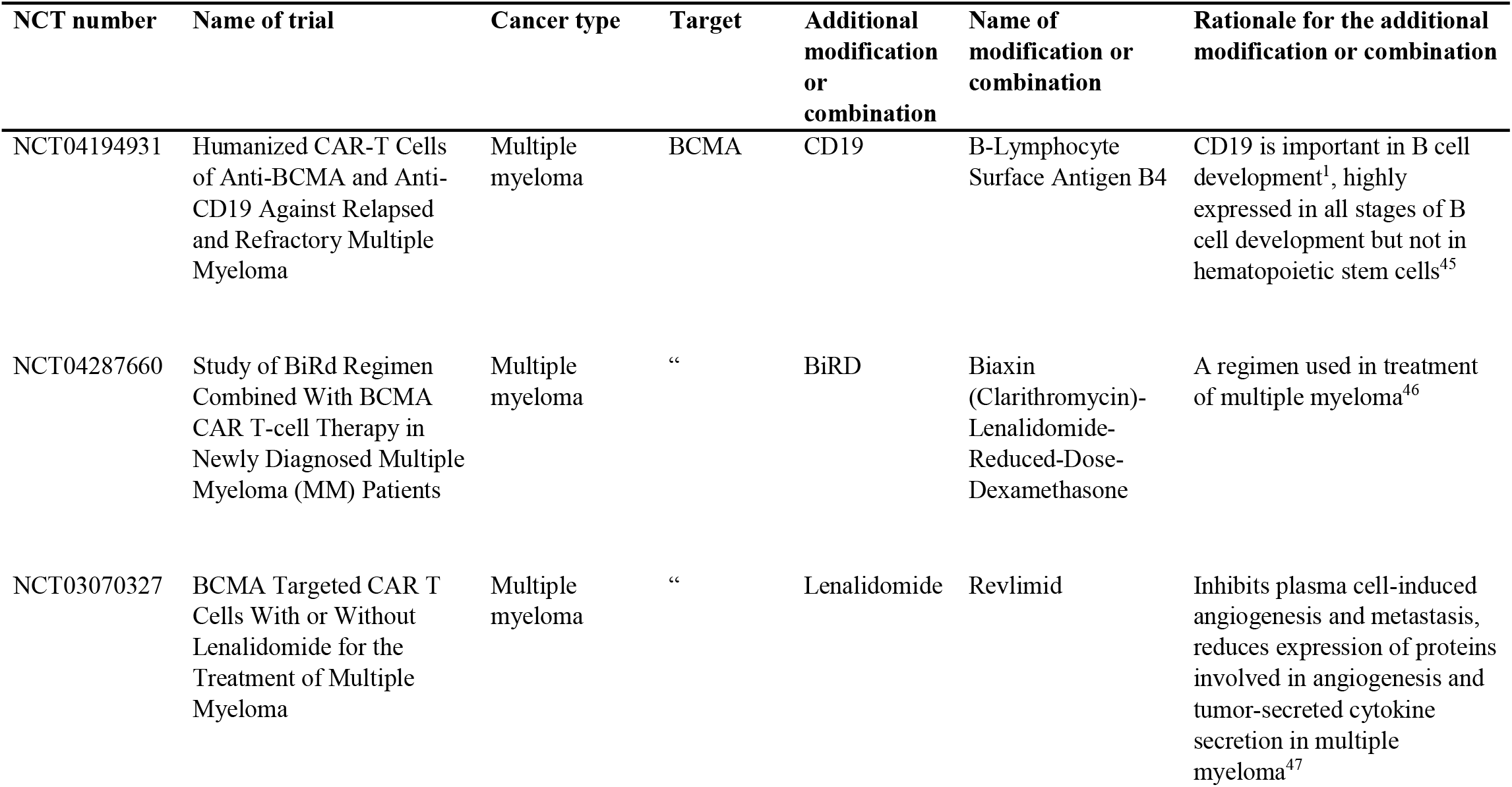
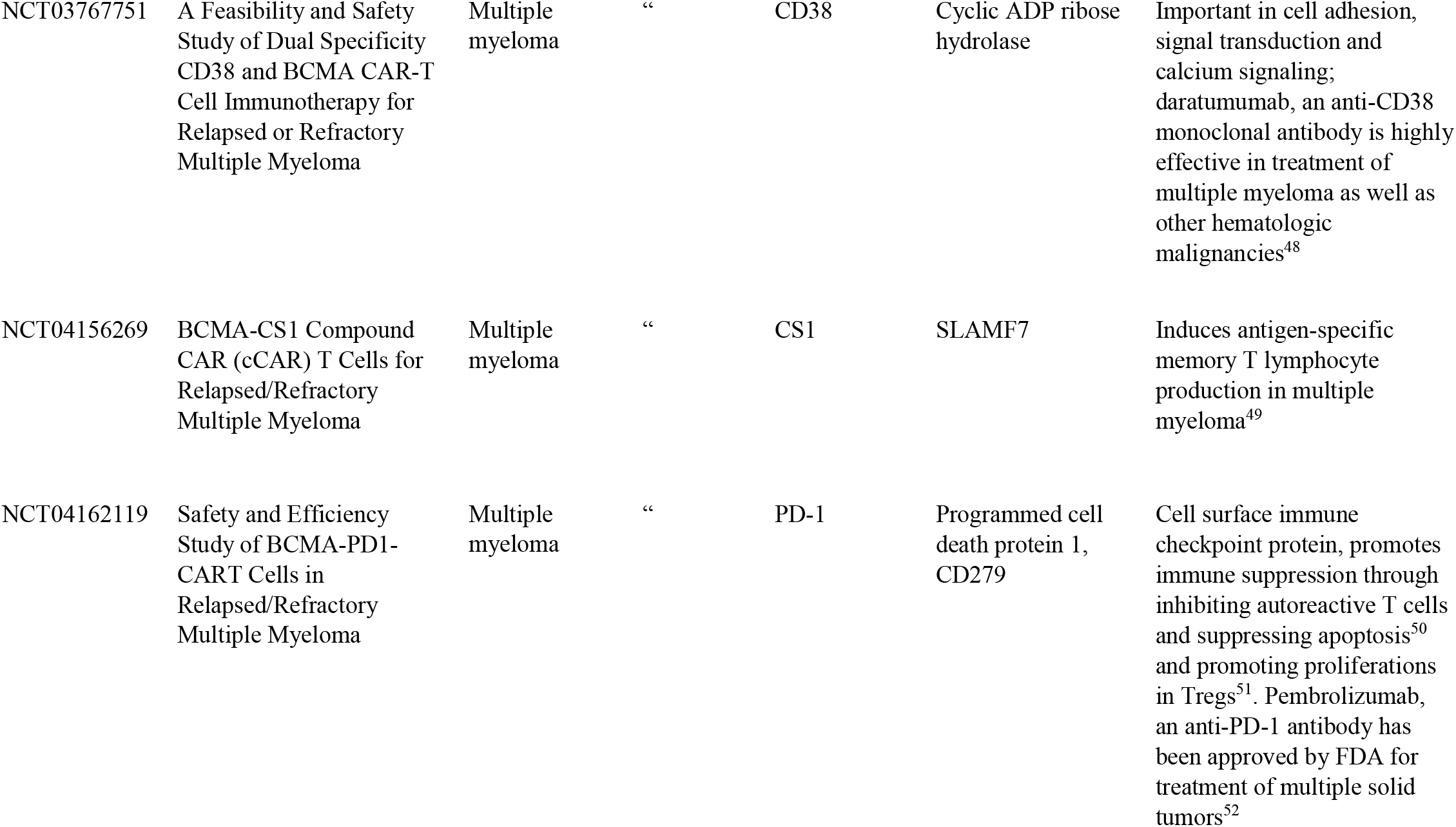
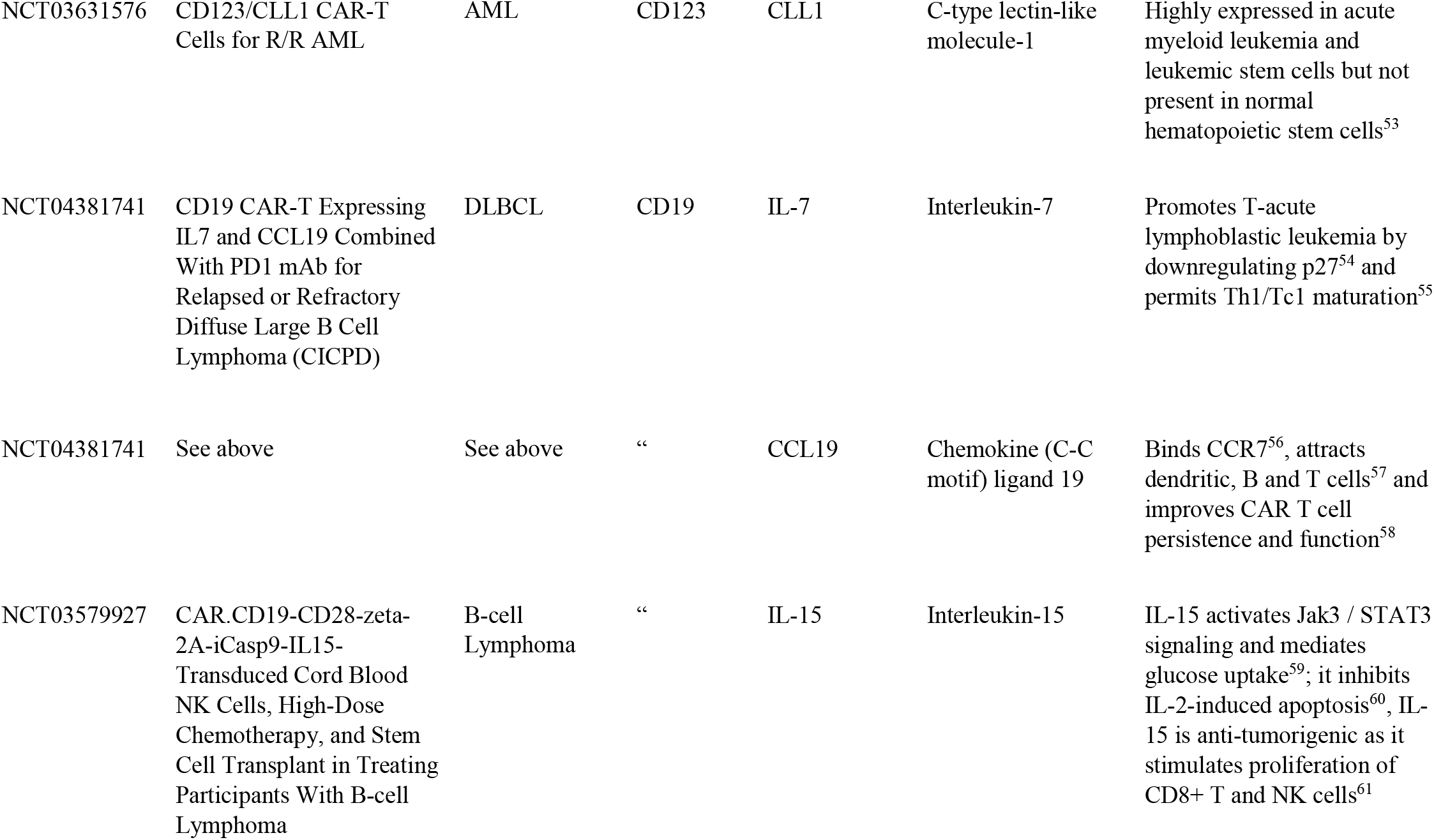
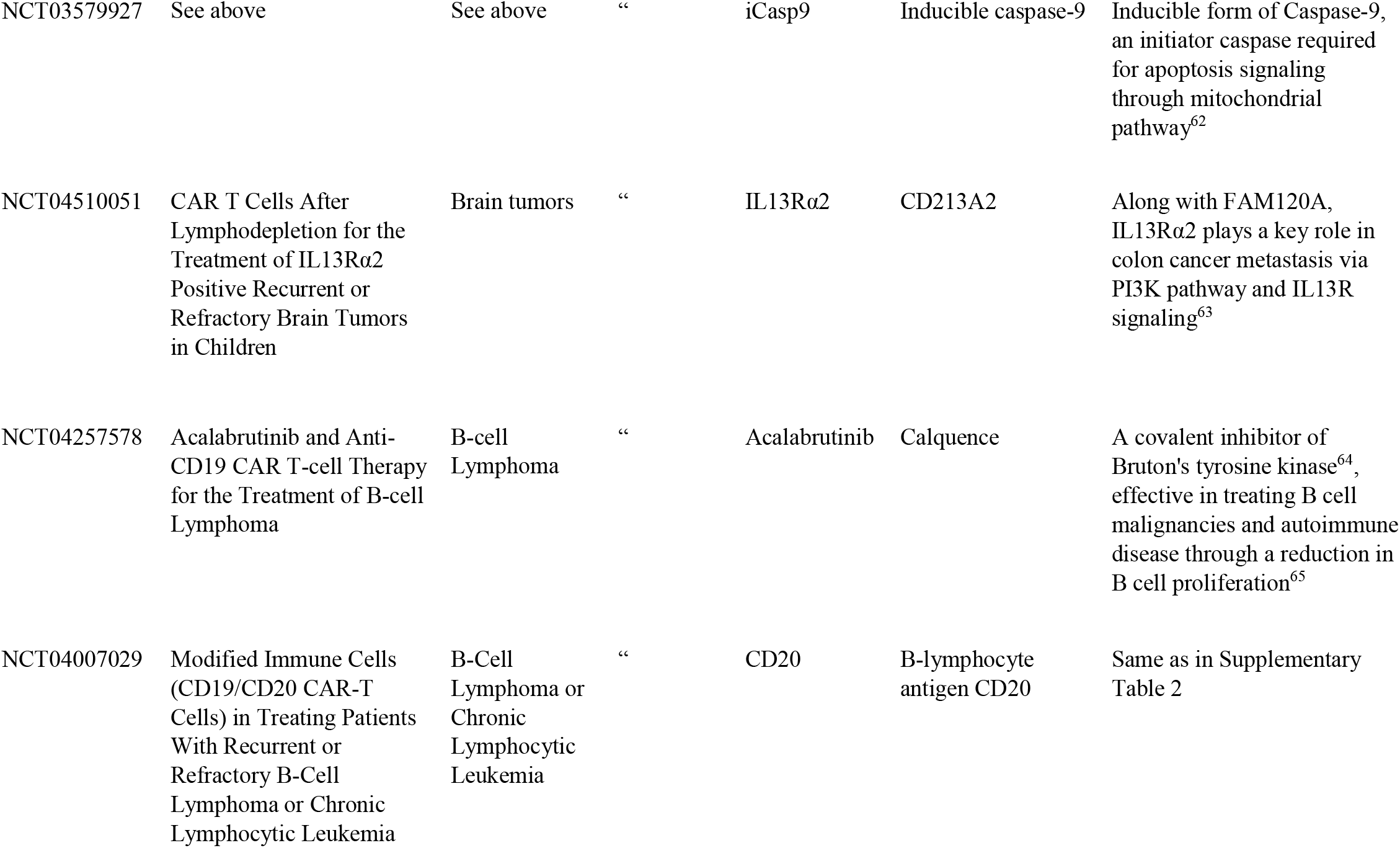
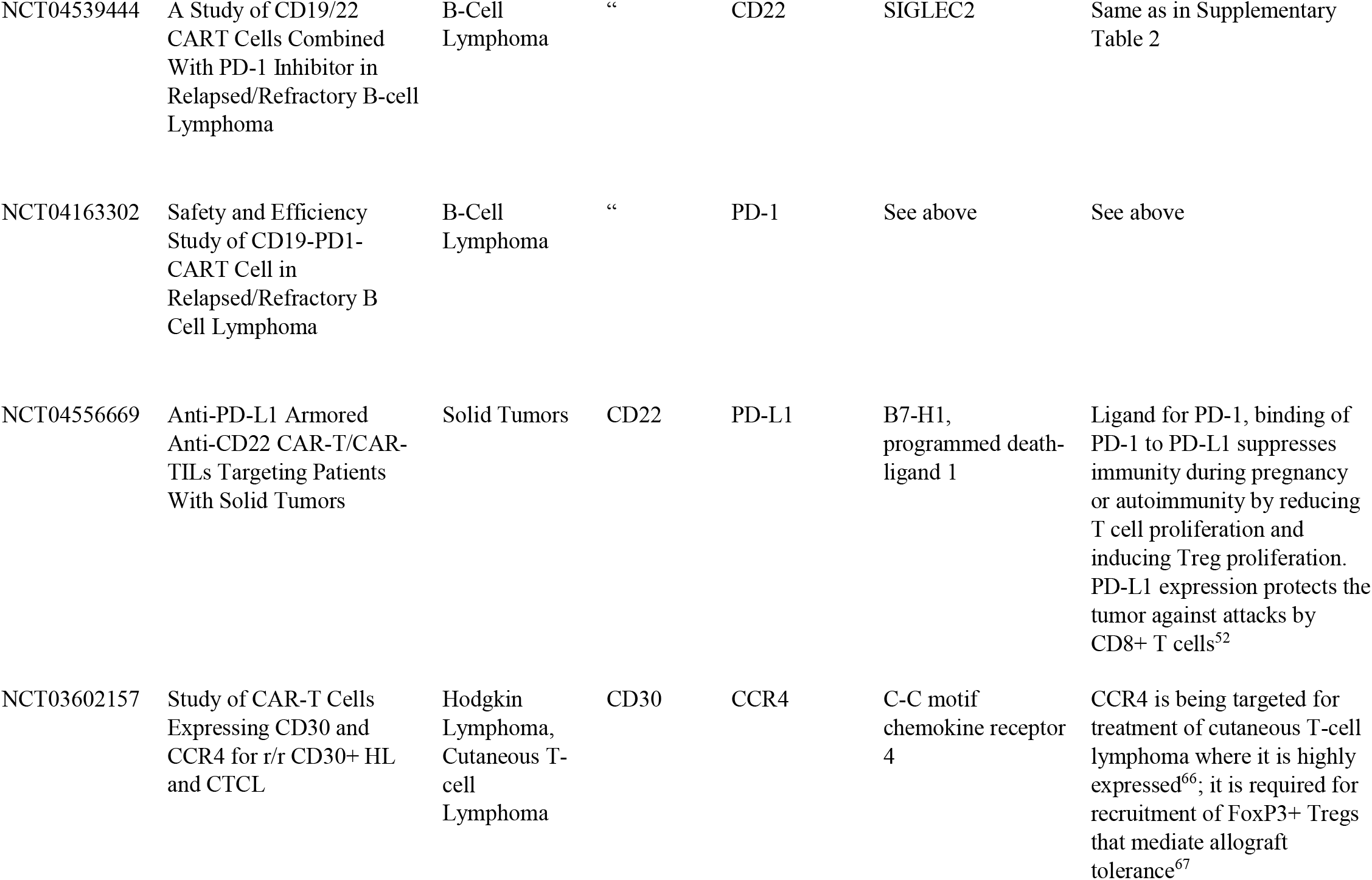
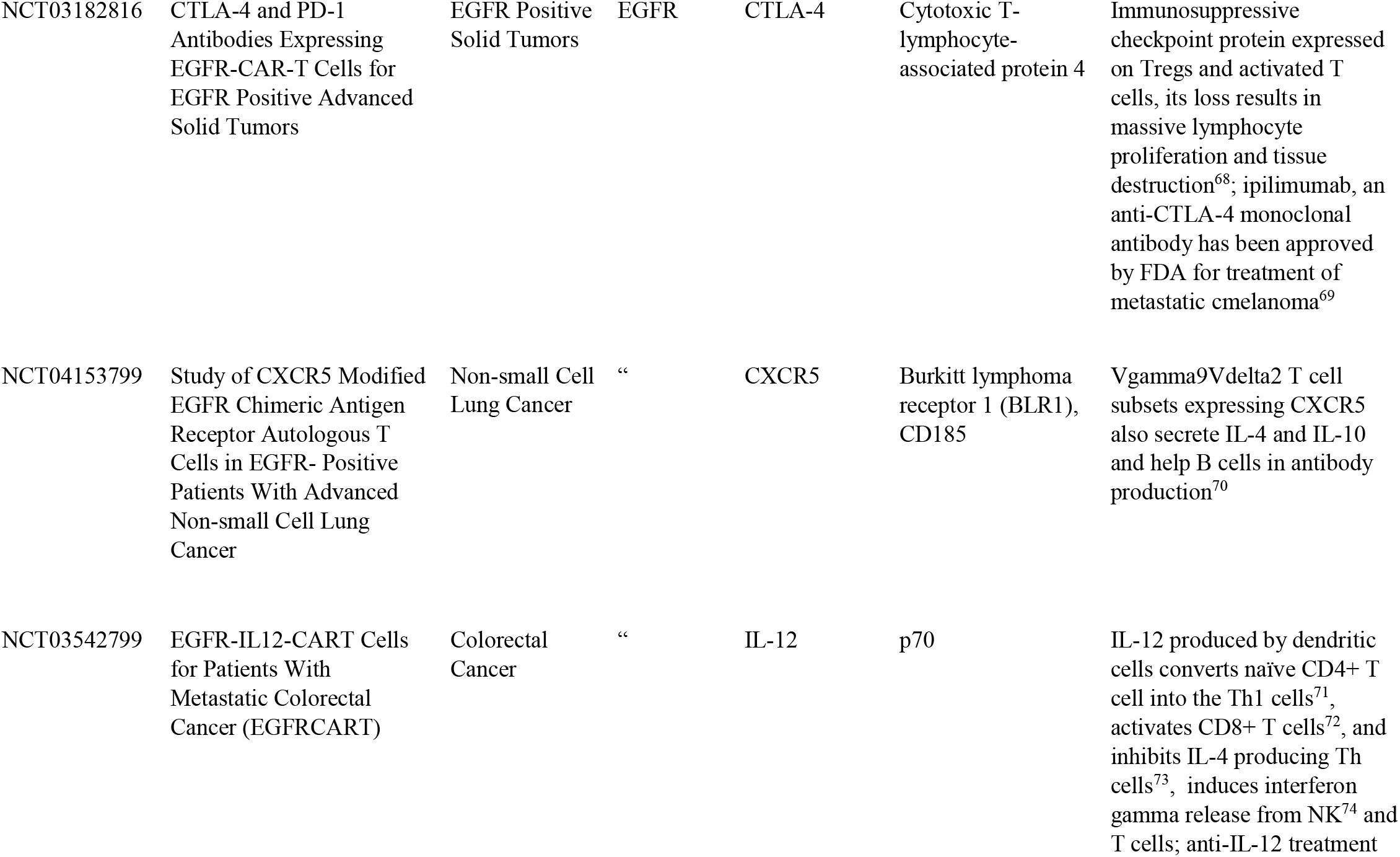
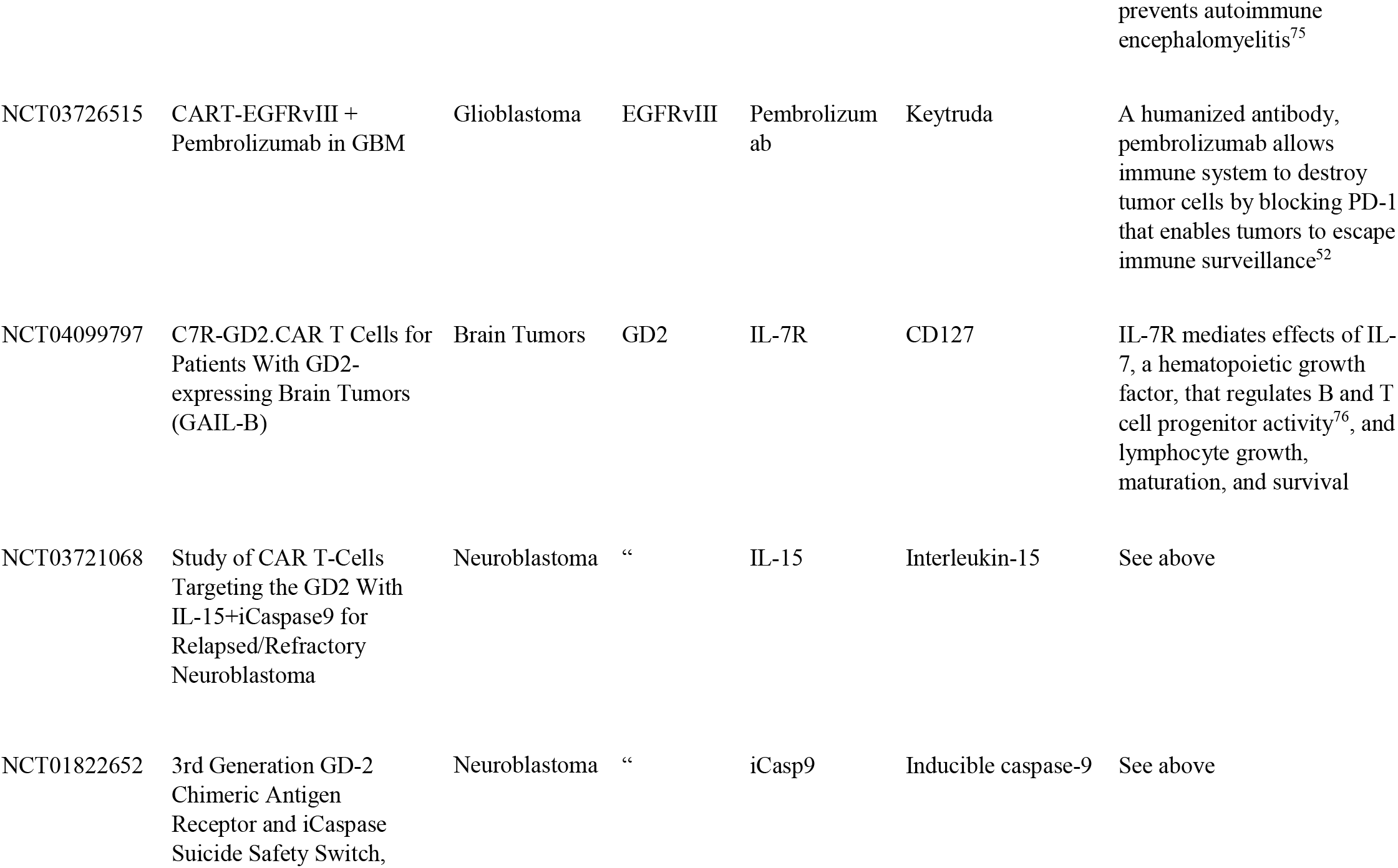
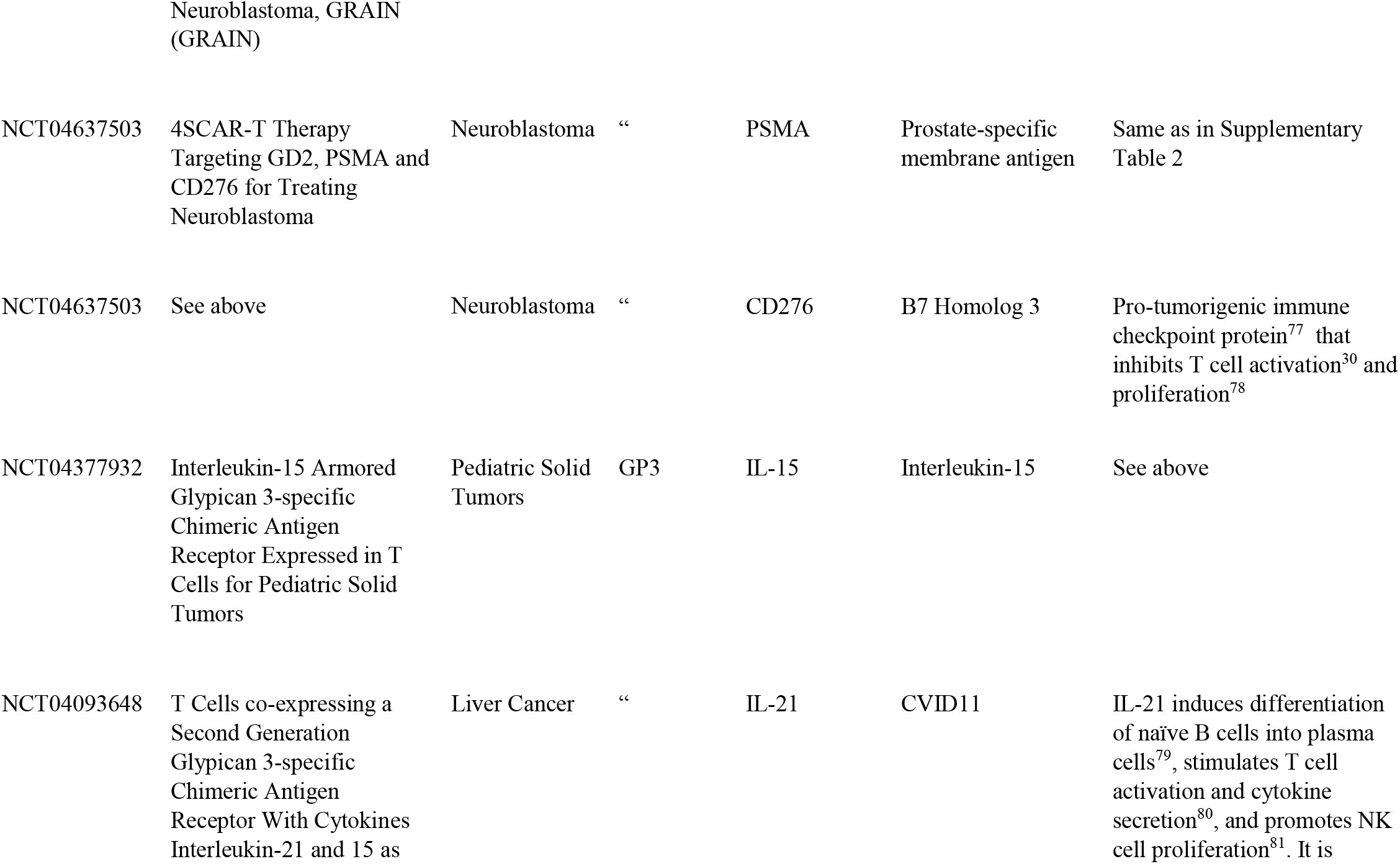
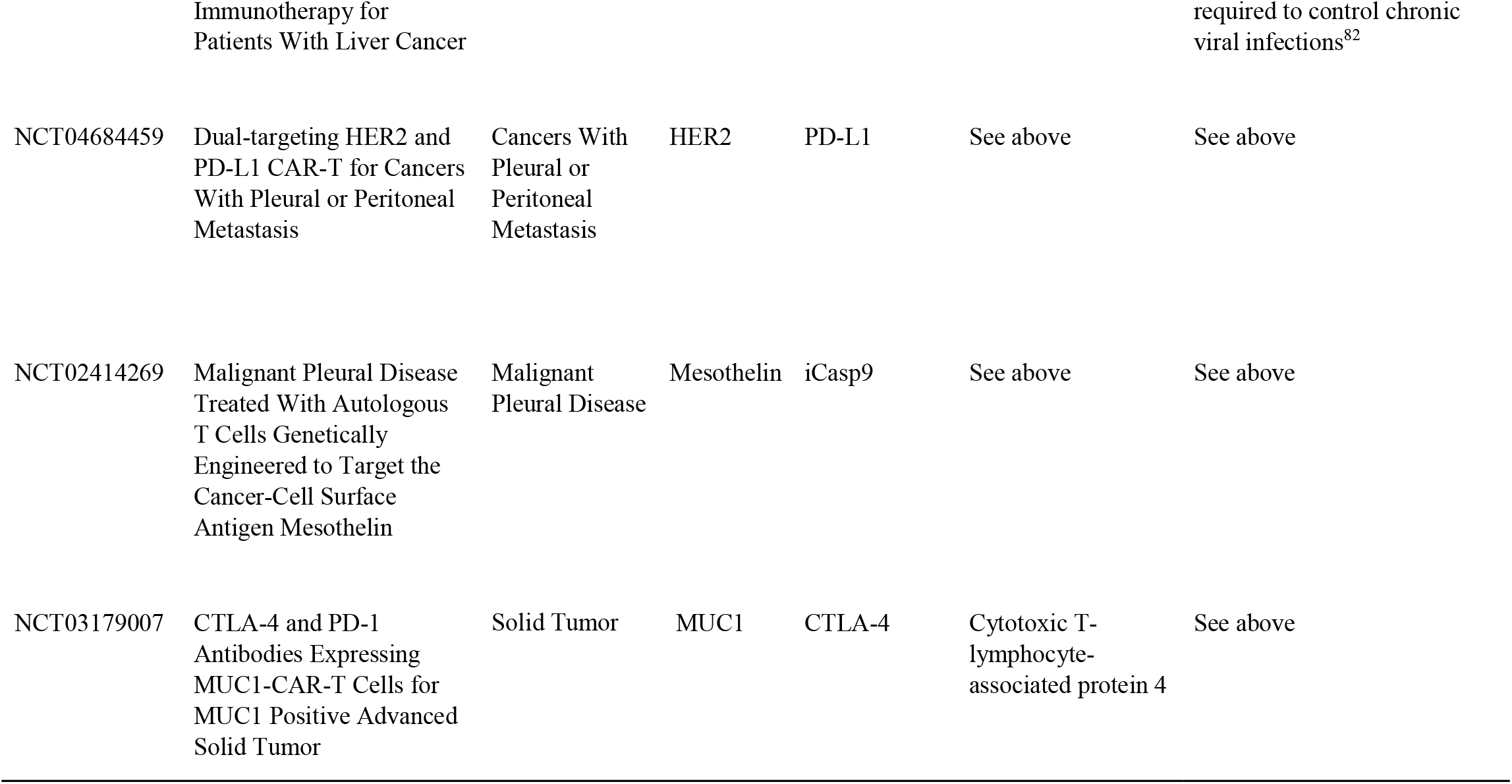
Additional modifications or compounds used to improve CARi efficacy in clinical trials.

**Supplementary Table 4.** Clinical trials targeting multiple CARs.

| **NCT number** | **Year** | **Targets** | **Condition** |
| --- | --- | --- | --- |
| NCT04430530 | 2020 | CD22/CD123/CD38/CD10/CD20 | CD19 negative B-cell Malignancies |
| NCT04429438 | 2020 | CD19/CD20/CD22/CD70/PSMA/CD13/CD79b/GD2 | B Cell Lymphoma |
| NCT04010877 | 2019 | CLL-1/CD33/CD123 | Acute Myeloid Leukemia |
| NCT03778346 | 2018 | Integrin β7/BCMA/CS1/CD38/CD138 | Multiple Myeloma |
| NCT03125577 | 2017 | CD19/CD20/CD22/CD30/CD38/CD70/CD123 | B-cell Malignancies |
| NCT03198052 | 2017 | HER2/Mesothelin/Lewis-Y/PSCA/MUC1/GPC3/AXL<br>/EGFR/B7-H3/Claudin18.2 | Lung Cancer |
| NCT03222674 | 2017 | Muc1/CLL1/CD33/CD38/CD56/CD123 | Acute Myeloid Leukemia |
| NCT03196414 | 2016 | CD138/BCMA/CD19 | Multiple Myeloma |

**Supplementary Table 5.**
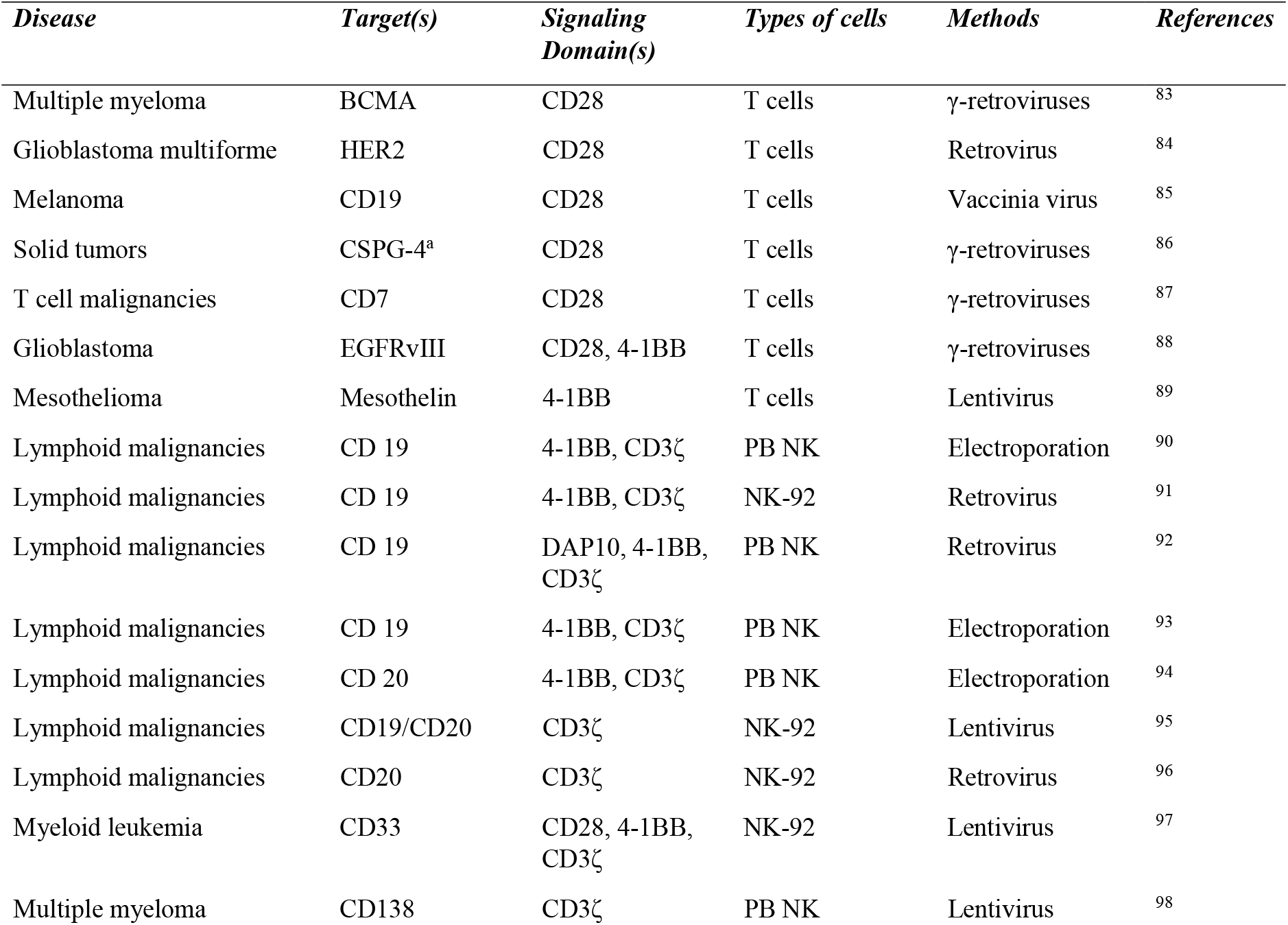
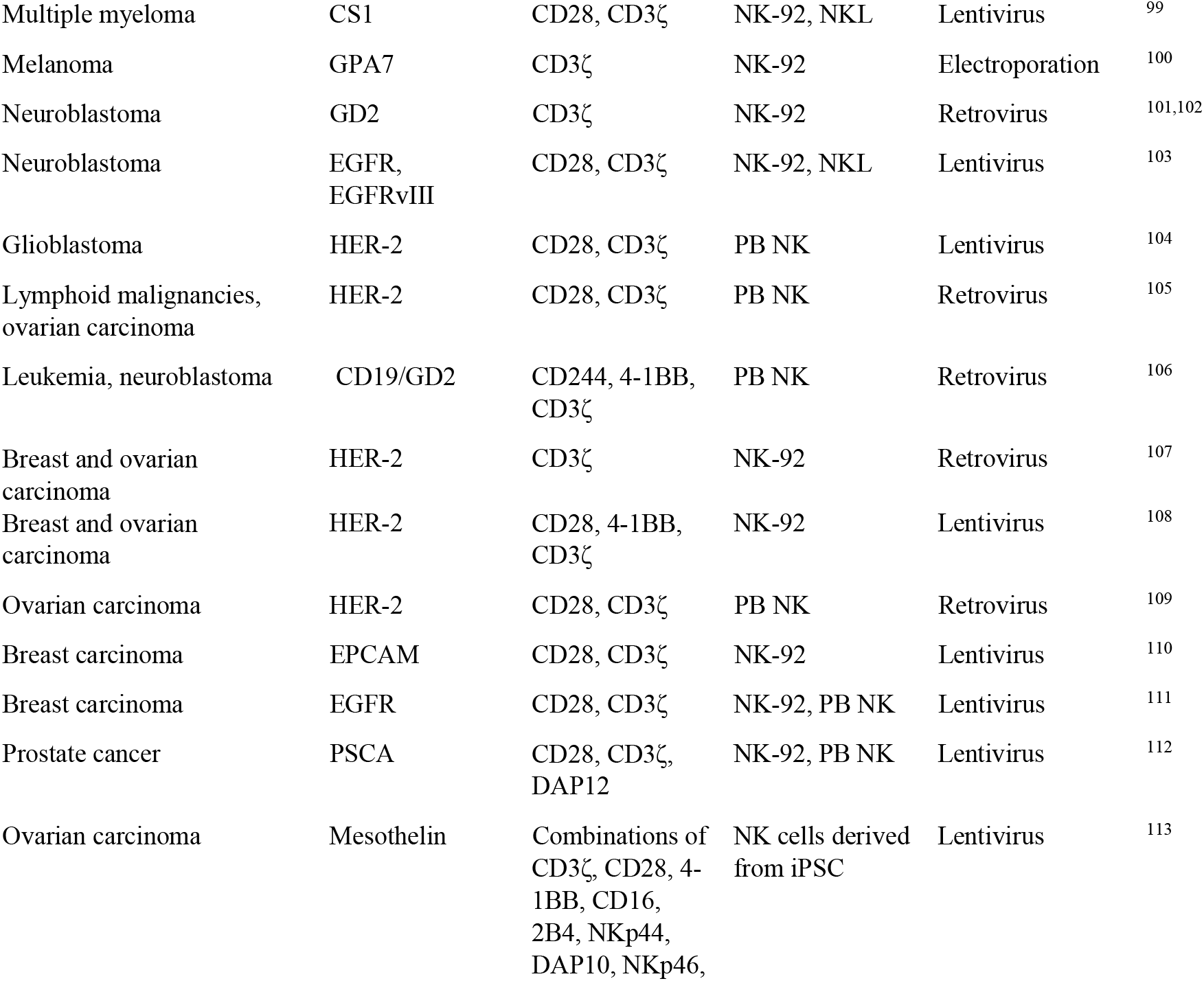
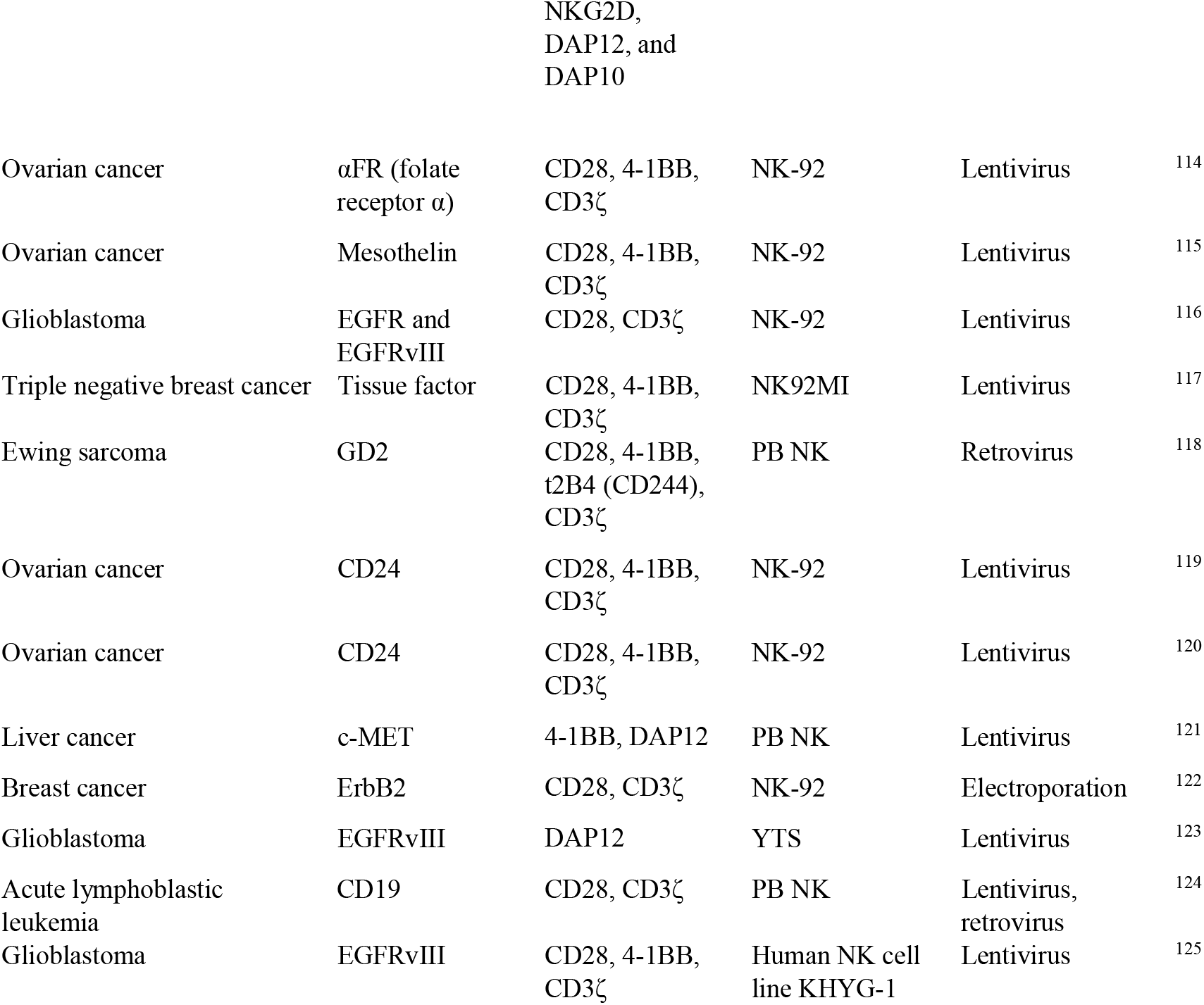
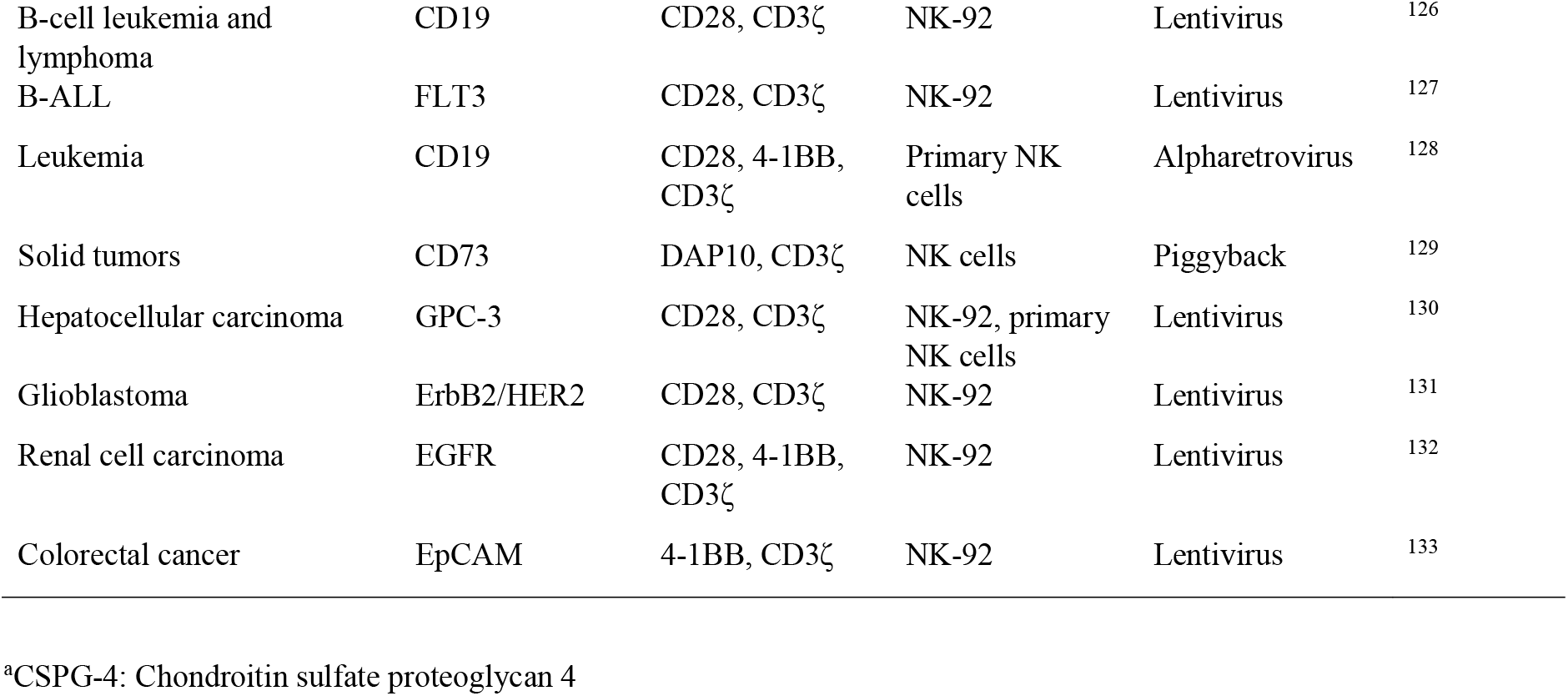
Representative preclinical studies of CAR T and CAR NK cell-based cancer immunotherapy.

**Supplementary Table 6.** Notable institutions performing CARi clinical trials^a^

| **Institution** | ***N*** |
| --- | --- |
| Zhejiang University Hospitals, Zhejiang, China | 30 |
| People's Liberation Army General Hospitals, Beijing, China | 25 |
| University of Pennsylvania | 22 |
| City of Hope National Medical Center | 19 |
| National Institutes of Health | 19 |
| Houston Methodist Hospital | 17 |
| University of Washington | 13 |
| Yanda Ludaopei Hospital, Hebei, China | 13 |
| University of North Carolina | 13 |
| 3rd Military Medical University, Chongqing, China | 12 |
| Memorial Sloan Kettering Cancer Center | 10 |
| Seattle Children's Hospital | 10 |
| Texas Children's Hospital | 8 |
| Tongji Medical College, Wuhan, Hubei, China | 8 |
| Xinqiao Hospital, Chongqing, China | 7 |
| PersonGen Biomedicine, Jiangsu, China | 7 |
| Soochow university, Jiangsu, China | 7 |
| University of Alabama at Birmingham | 7 |
| University of California, San Francisco | 5 |

<sup>a</sup>Only the lead institution is included whenever multiple institutions were involved. Closely related institutions such as Abramson Cancer Center and University of Pennsylvania are combined. *N* = number of clinical trials performed by the institution.

**Supplementary Table 7.** An overall comparison of the anti-CD19 CAR T cell products approved by the FDA^a^

|  | **Tisagenlecleucel** | **Axicabtagene ciloleucel** | **Brexucabtagene autoleucel** | **Lisocabtagene maraleucel** |
| --- | --- | --- | --- | --- |
| **Brand name** | Kymriah<sup>134</sup> | Yescarta<sup>134</sup> | Tecartus<sup>135</sup> | Breyanzi |
| **Approval date** | August, 2017<sup>135</sup>: FDA-approved for B-ALL, May, 2018<sup>136</sup>: FDA-approved for adult DLBCL | October, 2017: FDA-approved for DLBCL<sup>137,138</sup> | July, 2020<sup>139</sup>: FDA-approved for MCL | February, 2021: FDA-approved for DLBCL |
| **The firsts** | The first FDA-approved gene therapy<sup>140</sup> (along with Luxturna) | The first FDA-approved CAR-T cell product for large B cell lymphoma<sup>139</sup> | The first cell-based therapy for MCL, an aggressive type of B cell non-Hodgkin's lymphoma seen mostly in adults over 60 years of age with no proven therapy regimen<sup>141</sup> | The first CAR T cell therapy with the regenerative medicine advanced therapy designation |
| **Patients** | Adults with r/r DLBCL and young adults or children under 25 years of age<sup>142</sup> with r/r B-cell precursor ALL | Adults | Adults | Adults |
| **Developer** | Developed at the University of Pennsylvania and Children's Hospital of Pennsylvania; further developed and marketed by Novartis<sup>143</sup> | Kite Pharma<sup>144</sup> (acquired by Gilead in 2017) | Kite Pharma | Juno therapeutics (acquired by Celgene in 2018) |
| **Pivotal trial** | JULIET phase II trial<br>(NCT02445248) | ZUMA-1 (Phase I/II) | ZUMA-2 (Phase II)<sup>145</sup><br>(NCT02601313) | TRANSCEND-NHL-001<br>(Phase I)<sup>146</sup><br>(NCT02631044) |
| **Co-stimulatory**<br>**domain**<sup>147</sup> | 4-1BB | CD28 | CD28 | 4-1BB |
| **Patients reported to the FDA Adverse Event Reporting System** |  |  |  |  |
| **Patients with AEs** | 1,546 | 2,072 | 3 | 50 |
| **Patients with**<br>**serious AEs** | 1,461 (94.50%) | 1,972 (95.17%) | 2 | 50 |
| **Death rate<sup>b</sup>** | 435 (28.14%) | 332 (16.02%) | 1 (50%) | 9 (18%) |

<sup>a</sup>r/r = relapsed/refractory; nos = not otherwise specified; MCL = mantle cell lymphoma, AE = adverse event.
<sup>b</sup>Death rate as a percentage of serious adverse events.

